# Temporal and Regional Circular RNA profiling in a Tauopathy Mouse Model: Implications for Tau Pathology and Neurodegeneration

**DOI:** 10.64898/2026.06.18.733253

**Authors:** Mohammad Shahadat Hossain, Domenico Praticò

## Abstract

MicroRNAs (miRNA), are non-coding RNA that act as post-transcriptional regulators of gene expression in various organs including the brain where they play an important role in neurodegeneration. Circular RNAs are single-stranded, covalently closed loop RNA molecules recognized as upstream regulators of miRNA. Previous studies have shown that circRNAs are dysregulated in Alzheimer’s and other neurodegenerative diseases. However, a systematic, age-and region-specific circRNA atlas in primary tauopathy is lacking.

To this end, we performed comprehensive circRNA sequencing of hippocampal and cortical tissues from a model of human tauopathy, h-Tau mice, at 3, 6, and 12 months of age. We identified circRNA–miRNA sponging networks that target and regulate key tau disease-associated pathways, including kinases, phosphatases, histone deacetylase, glutamatergic and GABAergic synapse, and microglial efferocytosis.

Our study demonstrates an age- and region-specific circRNA landscape in the brain of a model of human tauopathy and identify candidate circRNA–miRNA–mRNA regulatory axes converging on core tau pathological processes. These findings support the novel hypothesis that specific circRNAs have the potential to be used as biomarkers and therapeutic targets against tau-driven neurodegeneration.

## 1. Introduction

Tauopathies, including Alzheimer’s disease (AD), Progressive supranuclear palsy (PSP), Corticobasal degeneration (CBD) and Pick’s disease, are disorders characterized by intraneuronal accumulation of hyperphosphorylated tau protein.(Nichols et al., 2022; Spillantini and Goedert, 2013) Tau normally stabilizes axonal microtubules, but its aberrant phosphorylation by kinases such as GSK3β, ERK1/2, JNK, and p38 MAPK at multiple epitopes (Thr181, Thr231, Ser202, Ser396, Ser422) causes its detachment, somatodendritic mislocalization, and aggregation into insoluble filaments that then become neurofibrillary tangles.(Hanger et al., 2009; Iqbal et al., 2016; Wang and Mandelkow, 2016; You et al., 2015) Tau pathology propagates in a circuit-dependent manner, originating in the entorhinal cortex and hippocampal CA1 before advancing outward through interconnected cortical networks as defined by the Braak staging framework,(Braak and Braak, 1991; de Calignon et al., 2012) with the hippocampus and cerebral cortex representing the two regions of greatest vulnerability.

Non-coding RNAs have emerged as critical post-transcriptional regulators of gene expression in the brain, with microRNAs (miRNAs), short ∼22-nucleotide regulatory RNAs that suppress target mRNA translation or promote degradation,(Bartel, 2009) playing a prominent role in neurodegeneration. Dysregulated miRNA profiles have been documented in AD brain tissue and biofluids.(Cha et al., 2019; Delay and Hebert, 2011; Hebert and De Strooper, 2009; Wang et al., 2009) Our group previously profiled miRNA expressions in the cortex and hippocampus of a mouse model of tauopathy, h-Tau mice, across disease progression, revealing distinct region- and age-specific miRNA signatures.(Lauretti et al., 2021)

Circular RNAs (circRNAs) covalently closed RNA molecules generated by back-splicing of pre-mRNA exons,(Memczak et al., 2013) are now recognized as major upstream regulators of miRNA. Their closed-loop architecture confers exceptional metabolic stability in post-mitotic neurons, and the brain expresses thousands of circRNA species in a cell type-specific fashion.(Rybak-Wolf et al., 2015; You et al., 2015) Critically, circRNAs function as competitive endogenous RNA (ceRNA) sponges, sequestering miRNAs through tandem miRNA response elements, and thereby de-repressing the cognate target mRNAs.(Hansen et al., 2013; Piwecka et al., 2017; Salmena et al., 2011)

Dysregulated circRNA profiles have been described in AD cortex(Dube et al., 2019) and Parkinson’s disease substantia nigra,(Gao et al., 2025; Hanan et al., 2020) but a systematic, age-and region-specific circRNA atlas in primary tauopathy, a condition where tau is the only pathologic feature, is lacking. To address this gap, we performed comprehensive circRNA sequencing of hippocampal and cortical tissues from h-Tau mice, a model of primary tauopathy, at 3, 6, and 12 months of age. We identified circRNA–miRNA sponging networks that regulate key disease-associated pathways, including tau phosphorylation and dephosphorylation,(Gong et al., 1994) epigenetic modifications,(Graff et al., 2012) glutamatergic and GABAergic synapses, and microglial efferocytosis.(Brown and Neher, 2014)

Our findings support the novel hypothesis that specific circRNAs have the potential to be used not only as biomarkers but also as therapeutic targets against tau-driven neurodegeneration.

## 2. Materials and Methods

### 2.1. Animals and tissue collection

Human tau (hTau) transgenic mice mouse (JAX #005491) was used in this study. Animals were grouped into three age cohorts: 3 months, 6 months, and 12 months, with n = 4 mice per age group and an equal number of males and females in each cohort. Mice were housed under standard laboratory conditions with ad libitum access to food and water and maintained on a 12 h light/dark cycle. All animal procedures were conducted in strict accordance with the National Institutes of Health (NIH) Guide for the Care and Use of Laboratory Animals and were reviewed and approved by the Institutional Animal Care and Use Committee (IACUC) of Temple University. All efforts were made to minimize animal suffering and to reduce the number of animals used. Mice were euthanized using carbon dioxide (CO₂) inhalation in an IACUC-approved euthanasia chamber. CO₂ was delivered from a compressed gas cylinder equipped with a regulator and flow meter set to a flow rate of 4 liters per minute (LPM), corresponding to a gradual fill rate consistent with institutional guidelines for adult mice. Following placement of the animal into the chamber, CO₂ flow was initiated and maintained until respiration ceased. Animals were observed for an additional 30–60 seconds after cessation of breathing to ensure complete euthanasia prior to removal from the chamber. Immediately after euthanasia, mice were perfused with ice-cold phosphate-buffered saline (PBS) to remove circulating blood. Brains were rapidly extracted, and the hippocampus and cerebral cortex were carefully dissected on ice. Dissected tissues were snap-frozen on dry ice, stored at −80 °C, and shipped on dry ice to Novogene for RNA isolation, circRNA sequencing, and downstream bioinformatics analyses performed in-house.

### 2.2. RNA isolation and quality control

Total RNA was isolated in-house by Novogene using their standard operating procedures for circRNA sequencing. RNA concentration was measured using a Qubit Fluorometer, RNA integrity was assessed using the Agilent 5400 system, and sample purity was evaluated by agarose gel electrophoresis in combination with Agilent analysis. All RNA samples passed Novogene’s quality control criteria for circRNA library preparation, with RNA integrity values ranging from 7.3 to 9.0, and were approved for downstream processing.

### 2.3. circRNA library preparation and sequencing

Circrna libraries were prepared in-house at Novogene using total RNA as input. Ribosomal RNA (rRNA) was removed using rRNA-specific probes, followed by ethanol precipitation. Linear RNAs were selectively digested with RNase R to enrich circular RNAs. The enriched RNA was fragmented, and first-strand cDNA synthesis was performed using random hexamer primers. Second-strand cDNA synthesis was carried out using dUTP instead of dTTP to maintain strand specificity. Libraries were generated through end repair, A-tailing, adapter ligation, size selection, PCR amplification, and purification. Library quality assessment included Qubit quantification and insert size evaluation. Final library concentrations were determined by quantitative real-time PCR (qRT-PCR). Qualified libraries were pooled and sequenced by Novogene on an Illumina sequencing platform to generate paired-end reads.

### 2.4. Raw data processing and read mapping

Base calling and conversion of raw fluorescence signals into FASTQ files were performed using Novogene’s sequencing pipeline. Quality control of raw reads was conducted using fastp (v0.23.1). Reads were filtered to remove: (1) adapter-contaminated reads; (2) reads containing >10% ambiguous nucleotides (N); and (3) reads with >50% low-quality bases (Phred score <5). Clean reads were aligned to the reference genome by Novogene using HISAT2 (v2.2.1). Mapping results were used to support accurate detection of back-splice junction reads required for circRNA identification.

### 2.5. circRNA identification, quantification, and differential expression analysis

Circular RNAs were identified by Novogene using both Find_circ v1.2 (Memczak et al., 2013) and CIRI2 v2.0.6 (Gao et al., 2015). circRNA expression levels were quantified based on back-splice junction read counts and normalized using Transcripts Per Million (TPM). Differential expression analysis was performed by Novogene using DESeq2 v1.42.0 (Love et al., 2014)for datasets with biological replicates. For comparisons lacking biological replicates, edgeR v4.0.16 (Robinson et al., 2010)with the trimmed mean of M values (TMM) normalization was applied. circRNAs with adjusted *p* values < 0.05 were considered significantly differentially expressed.

### 2.6. Selection of circRNAs for downstream analysis

Based on differential expression results, four circRNAs were selected for downstream analysis and functional characterization. Two circRNAs were selected from each brain region (hippocampus and cerebral cortex), including one upregulated and one downregulated circRNA per region. Selection criteria included statistical significance, magnitude of expression change, and the biological relevance of parental genes, with emphasis on genes implicated in neurodegeneration and synaptic dysfunction, as supported by functional enrichment analysis and literature evidence.

### 2.7. circRNA – miRNA –mRNA interaction prediction

circRNA–miRNA interactions were predicted using miRanda softwaere and Putative miRNA–mRNA interactions were predicted using miRDB. Selected miRNA–mRNA interactions were incorporated into downstream functional analyses and circRNA–miRNA–mRNA regulatory network construction via Cytoscape v3.10.4 (Shannon et al., 2003) visualization.

### 2.8. Functional enrichment analysis

Functional enrichment analyses of the selected circRNA host genes and miRNA target genes were performed using DAVID Bioinformatics Resources. Enrichment analysis was conducted for Biological Process, Molecular Function, Cellular Component categories, and KEGG pathway. GO and KEGG terms with *p* values < 0.05 were considered significantly enriched and were used for functional interpretation related to neuronal function, synaptic signaling, and neurodegenerative pathways.

## 3. Results

### 3.1. Region and age-Dependent differential Expression of circRNAs in tauopathy

To characterize circRNA expression dynamics during brain aging, we first applied unsupervised hierarchical clustering to all detected circRNAs across six experimental groups encompassing two brain regions (cortex and hippocampus) and three age points (3M, 6M, and 12M). The resulting heatmap (Figure 1a) revealed two dominant dendrogram branches that distinctly separated samples by brain region rather than age, with biological replicates clustering tightly within each condition. All cortex samples (3M, 6M, and 12M) were clustered together and clearly segregated from hippocampus samples (3M, 6M, and 12M), indicating that circRNA expression programs are fundamentally region-specific and establish a strong cortex-versus-hippocampus identity in the circRNA transcriptome.

**Figure 1.**
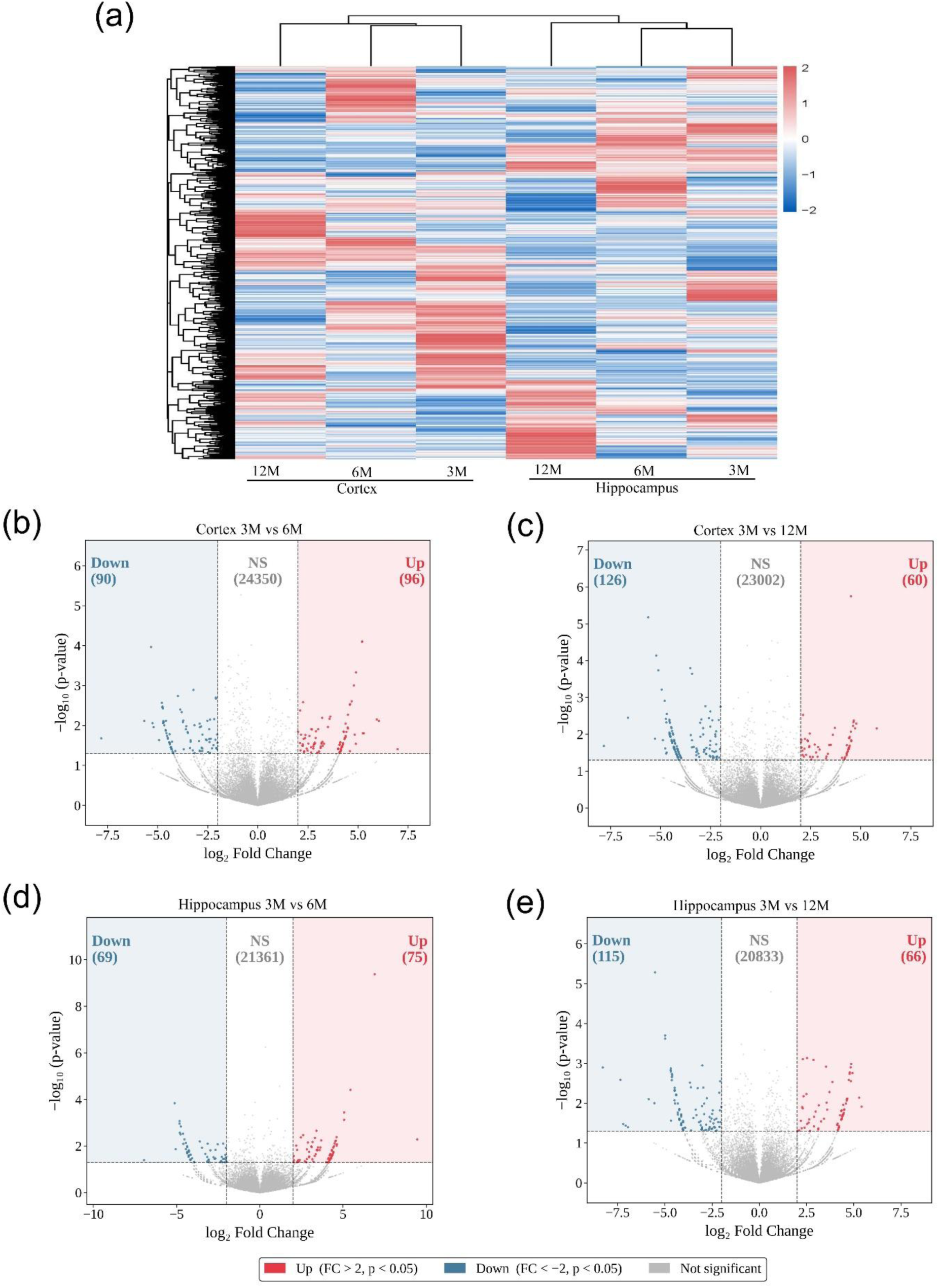
Differential expression of circular RNAs (circRNAs) in the cortex and hippocampus across disease progression in a tauopathy mouse model. (a) Hierarchical clustering heatmap depicting the expression profiles of differentially expressed circRNAs across three time points (3M, 6M, and 12M) in the cortex and hippocampus. Each row represents an individual circRNA and each column represents a sample group. Color scale indicates z-score normalized expression values (red: upregulated; blue: downregulated). (b) Volcano plot illustrating differentially expressed circRNAs in the cortex between 3M and 6M. (c) Volcano plot of differentially expressed circRNAs in the cortex between 3M and 12M. Significantly upregulated: n = 60; downregulated: n = 126; NS: n = 23,002. (d) Volcano plot of differentially expressed circRNAs in the hippocampus between 3M and 6M. Significantly upregulated: n = 75; downregulated: n = 69; NS: n = 21,361. (e) Volcano plot of differentially expressed circRNAs in the hippocampus between 3M and 12M. Significantly upregulated: n = 66; downregulated: n = 115; NS: n = 20,833. Red dots indicate significantly upregulated circRNAs (FC > 2, p < 0.05; n = 96), blue dots indicate significantly downregulated circRNAs (FC < −2, p < 0.05; n = 90), and gray dots represent non-significant (NS) circRNAs (n = 24,350).

To further interrogate age-associated circRNA changes within each region, we conducted pairwise differential expression analyses comparing 3M to later time points (6M and 12M), using volcano plots with thresholds of *p* < 0.05 and |log₂ fold change| > 2 applied to all detected circRNAs passing expression filtering (Figure 1b–1e). In the cortex, comparison of 3M versus 6M identified 96 upregulated and 90 downregulated circRNAs, with the majority of transcripts remaining unchanged (24,350 total; Figure 1b). By contrast, the 3M versus 12M comparison revealed a marked shift toward repression, with only 60 circRNAs upregulated and 126 downregulated (Figure 1c), consistent with the heatmap pattern in which Cortex12M samples diverged more strongly from the 3M baseline and suggesting progressive, age-associated transcriptional suppression. A similar pattern was observed in the hippocampus. The 3M versus 6M comparison identified 75 upregulated and 69 downregulated circRNAs from a background of 21,361 unchanged transcripts (Figure 1d). At 12M, the number of downregulated circRNAs increased to 115, while 66 circRNAs were upregulated from 20,833 unchanged transcripts (Figure 1e). Notably, although hippocampal samples retained strong region-specific clustering, within-region divergence from the 3M baseline increased with age, mirroring trends observed in the cortex.

### 3.2. Identification of candidate circRNAs

To identify circRNAs exhibiting sustained and reproducible dysregulation across aging, we intersected the significantly altered circRNAs from both age comparisons (3M vs 6M, and 3M vs 12M) using Venn diagram analysis (**Figures 2a and 3a**). This strategy enabled the identification of a temporally stable subset of circRNAs consistently dysregulated relative to the 3M baseline. In the cortex, 134 circRNAs were uniquely altered at 6M and 121 at 12M, while 30 circRNAs overlapped across both comparisons, comprising 10 upregulated and 20 downregulated transcripts. In the hippocampus, 123 and 160 circRNAs were specific to the 6M and 12M comparisons, respectively, with 21 circRNAs persistently dysregulated across both time points (9 upregulated and 12 downregulated). These shared circRNAs, representing a stable core of age-associated changes, were selected for downstream functional analyses. Consistent with their classification, bar graph visualization of log₂ fold-change values relative to 3M confirmed coherent and sustained expression trajectories for the 30 cortical and 21 hippocampal circRNAs at both 6M and 12M (**Figures 2b and 3b**).

**Figure 2.**
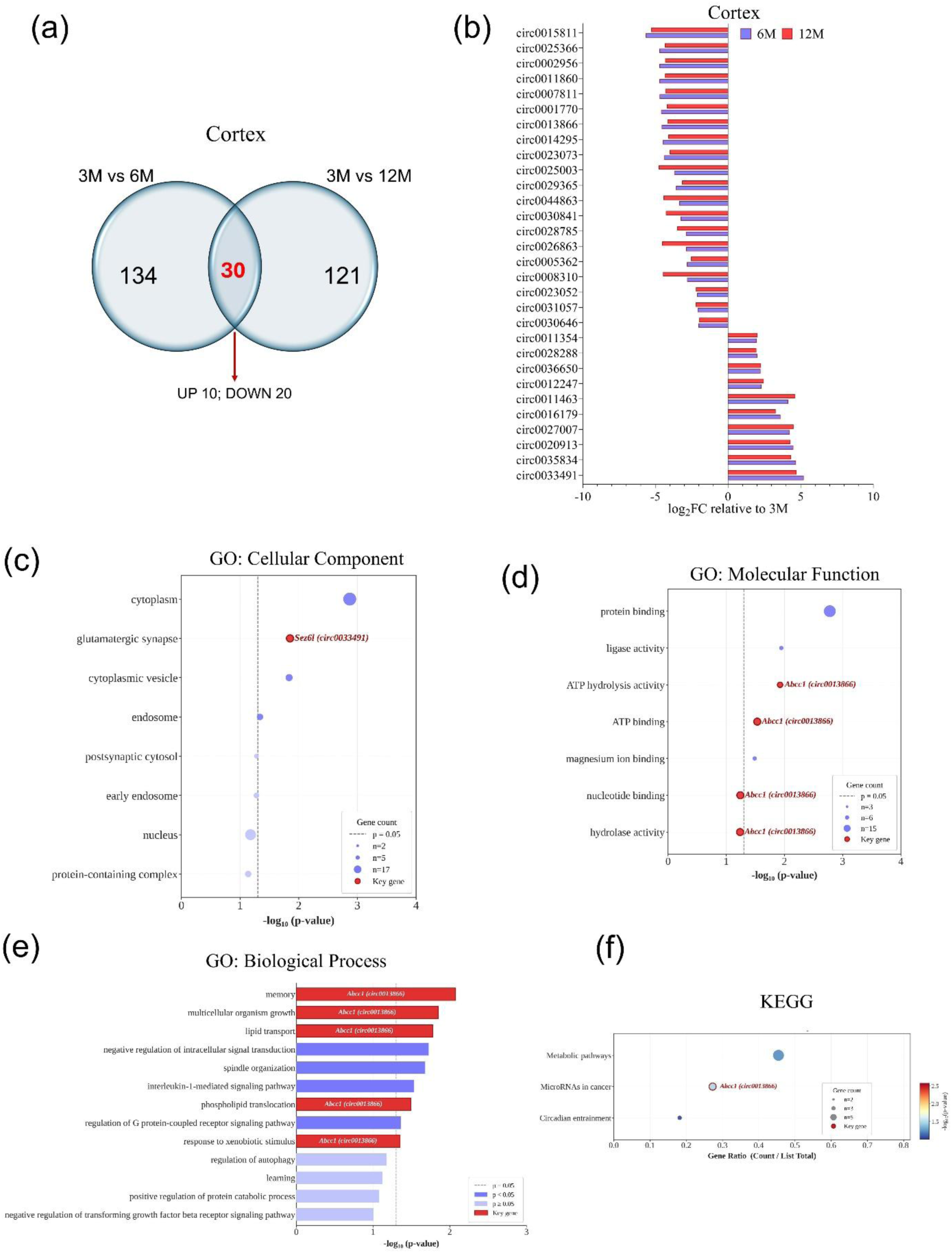
**Identification and functional annotation of consistently dysregulated circRNAs in the cortex**.(a) Venn diagram showing the overlap between differentially expressed circRNAs in the cortex at 3M vs. 6M (n = 134) and 3M vs. 12M (n = 121). Thirty circRNAs were commonly dysregulated across both comparisons, of which 10 were upregulated and 20 were downregulated. (b) Bar graph displaying the log₂ fold change (relative to 3M) of the top consistently differentially expressed circRNAs in the cortex at 6M (blue) and 12M (red). (c) Gene Ontology (GO) Cellular Component enrichment analysis of host genes associated with consistently dysregulated cortical circRNAs. Dot size reflects gene count and color intensity reflects statistical significance (−log₁₀ p-value). The key gene Sez6l (circ0033491) is annotated. (d) GO Molecular Function enrichment analysis. The key gene Abcc1 (circ0013866) is highlighted across multiple enriched terms. (e) GO Biological Process enrichment analysis, displaying bar lengths proportional to −log₁₀(p-value). Bars are color-coded by significance level (p < 0.01, p < 0.05, p ≥ 0.05) with key genes annotated. (f) Kyoto Encyclopedia of Genes and Genomes (KEGG) pathway enrichment analysis. Dot size represents gene count and color indicates −log₁₀(p-value). The key gene Abcc1 (circ0013866) is highlighted in enriched pathways. Dashed line indicates p = 0.05 significance threshold.

**Figure 3.**
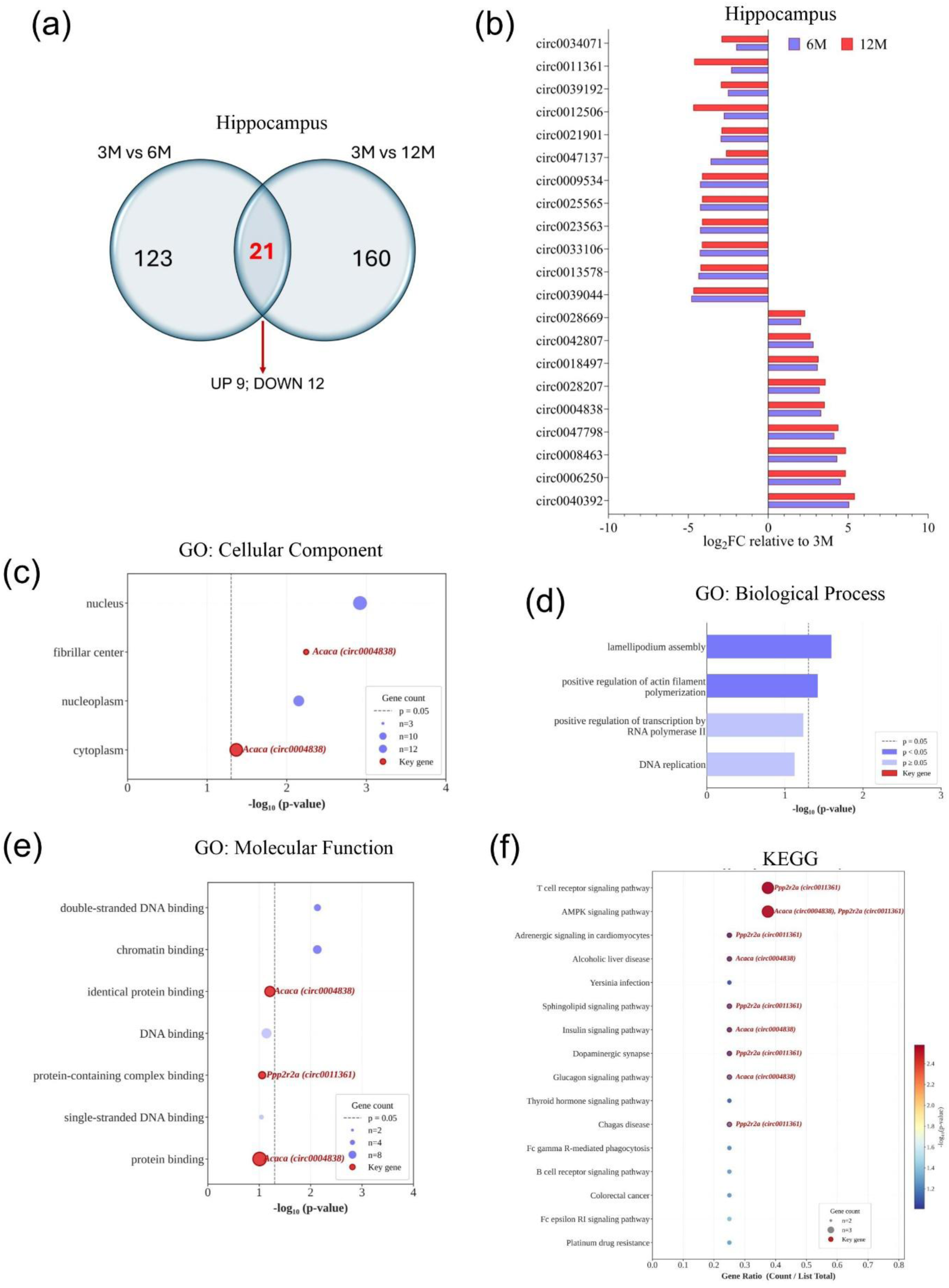
**Identification and functional annotation of consistently dysregulated circRNAs in the hippocampus**.(a) Venn diagram illustrating the overlap between differentially expressed circRNAs in the hippocampus at 3M vs. 6M (n = 123) and 3M vs. 12M (n = 160). Twenty-one circRNAs were consistently dysregulated, comprising 9 upregulated and 12 downregulated. (b) Bar graph showing the log₂ fold change (relative to 3M) of the top consistently differentially expressed circRNAs in the hippocampus at 6M (blue) and 12M (red). (c) GO Cellular Component enrichment analysis of host genes for consistently dysregulated hippocampal circRNAs. The key gene *Acaca* (circ0004838) is annotated in the cytoplasm and fibrillar center terms. Dot size indicates gene count and color reflects −log₁₀(p-value). (d) GO Biological Process enrichment analysis, bar colors indicate significance thresholds (p < 0.01, p < 0.05, p ≥ 0.05), with key genes annotated. (e) GO Molecular Function enrichment analysis. Key genes *Acaca* (circ0004838) and *Ppp2r2a* (circ0011361) are highlighted across terms. (f) KEGG pathway enrichment analysis of consistently dysregulated hippocampal circRNA host genes. Key genes *Ppp2r2a* (circ0011361) and Acaca (circ0004838) are highlighted in enriched pathways . Dot size reflects gene count; color indicates −log₁₀(p-value). Dashed line indicates p = 0.05.

To interpret the biological significance of the 30 consistently dysregulated cortical, functional enrichment analyses were performed on their parent genes using DAVID Bioinformatics Resources platform (**Supplementary information**, **S1 and S2**). GO Cellular Component (CC) analysis highlighted the cytoplasm (p = 0.0014) and glutamatergic synapses (p = 0.014), with *Sez6l* (circ0033491) specifically localizing to the glutamatergic synapses (**Figure 2c**), directly relevant to synaptic plasticity and memory. GO Molecular Function (MF) enrichment included protein binding (p = 0.0017) and ATP hydrolysis activity (p = 0.012) (**Figure 2d**). GO Biological Process (BP) analysis identified significant enrichment in memory (p = 0.0084), lipid transport (p = 0.017), multicellular organism growth (p = 0.014), and regulation of G protein-coupled receptor signaling (p = 0.043) (**Figure 2e**). *Abcc1*, the parent gene of circ0013866, was implicated in memory, lipid transport, phospholipid translocation, and xenobiotic response, suggesting its age-dependent downregulation may impair lipid homeostasis and cognitive function. KEGG analysis further identified enrichment in the MicroRNAs in cancer pathway (p = 0.034), implicating shared regulatory mechanisms between aging and oncogenic networks (**Figure 2f**).

Functional enrichment of parent genes for the 21 shared hippocampal circRNAs revealed distinct biological themes (**Supplementary information, S1 and S3**). GO CC analysis demonstrated enrichment in the nucleus (p = 0.0012) and fibrillar center (p = 0.0057), with *Acaca* contributing to nuclear compartment functions (**Figure 3c**). GO BP analysis identified enrichment in lamellipodium assembly (p = 0.025) and positive regulation of actin filament polymerization (p = 0.038), both critical for synaptic structural remodeling (**Figure 3d**). GO MF identified double-stranded DNA binding (p = 0.0073), chromatin binding (p = 0.0074), with *Ppp2r2a* associated with protein-containing complex binding (**Figure 3e**). KEGG pathway analysis revealed the most extensive functional associations, with the strongest enrichments in T cell receptor signaling (p = 0.0027, Ppp2r2a) and AMPK signaling (p = 0.0029, both *Acaca* and *Ppp2r2a*) a master regulator of cellular energy homeostasis critically implicated in aging and neurodegeneration (**Figure 3f**). Additional enriched pathways included insulin signaling (p = 0.089, *Acaca*), glucagon signaling (p = 0.067, *Acaca*), and dopaminergic synapse (p = 0.087, *Ppp2r2a*). The co-enrichment of both key hippocampal candidates in the AMPK pathway underscores their potential as coordinators of hippocampal energy homeostasis during aging.

Based on the persistence of dysregulation, magnitude of expression change, and functional relevance to neurodegeneration and tau pathology, four circRNAs were prioritized for further investigation. Among upregulated circRNAs, circ0033491 (*Sez6l*) was selected from the cortex and circ0004838 (*Acaca*) from the hippocampus. Circ0033491 showed robust and sustained induction with aging, reaching log₂FC ∼5.21 at 6M and 4.72 at 12M (**Figure 2b**), consistent with its clustering pattern in the heatmap indicating progressive cortical upregulation. Circ0004838 also demonstrated stable hippocampal upregulation, with log₂FC ∼3.31 at 6M and 3.53 at 12M (**Figure 3b**). Among downregulated circRNAs, circ0013866 (*Abcc1*) in the cortex exhibited consistent and substantial suppression (log₂FC ∼ −4.56 at 6M and −4.17 at 12M) (**Figure 2c**), aligning with its predicted involvement in lipid transport and neuronal homeostasis. Strikingly, circ0011361 (*Ppp2r2a*) in the hippocampus displayed an accelerated age-dependent decline, with log₂FC shifting from −2.32 at 6M to −4.63 at 12M (**Figure 3c**), mirroring the progressive suppression observed in heatmap and volcano plot analyses. Together, these expression trajectories and functional annotations highlight these four circRNAs as strong candidates for mediating age-associated molecular changes relevant to synaptic dysfunction, metabolic stress, and tau-related neurodegenerative processes.

### 3.3. Prediction of circRNA–miRNA-mRNA Network

CircRNAs function as competing endogenous RNAs (ceRNAs) by sequestering microRNAs (miRNAs) through direct base-pairing, thereby relieving miRNA-mediated suppression of downstream target mRNAs. To elucidate the regulatory potential of the four key aging-associated circRNAs identified in our expression analyses, we performed computational prediction of miRNA binding sites for circ0033491 (*Sez6l*) and circ0013866 (*Abcc1*) in the cortex, and circ0004838 (*Acaca*) and circ0011361 (*Ppp2r2a*) in the hippocampus. For downstream analyses, five miRNAs per circRNA were selected according to the following criteria:(1) Minimum free energy (MFE) ≤ −20 kcal/mol, with lower energy values indicating more thermodynamically stable circRNA–miRNA interactions; (2) Target binding score ≥ 150, reflecting a higher likelihood of true and strong targeting; (3) Lowest energy binding site among all predicted interactions for each circRNA–miRNA pair; and (4) Binding position and combinatorial binding sites, with preference given to miRNAs exhibiting multiple combinatorial binding positions on the same circRNA (**Table 1**). When multiple combinatorial binding sites were present, miRNAs with the highest binding score and most stable energy values were prioritized; miRNAs lacking combinatorial binding sites were deprioritized. Based on these criteria, the top five miRNAs with the strongest and most stable predicted interactions were selected for each circRNA and used for downstream functional and regulatory network analyses. More negative energy values and higher binding scores indicate stronger thermodynamic stability of the circRNA–miRNA duplex.

**Table 1.** Predicted miRNA binding interactions for the four key aging-associated circRNAs, ranked by total binding energy within each circRNA.

| circRNA | miRNA | Total Energy (kcal/mol) | Highest Binding Score | Lowest Energy (kcal/mol) | Binding Position(s) | miRNA Length | circRNA Length |
| --- | --- | --- | --- | --- | --- | --- | --- |
| circ0033491<br>(*Sez6l*) | miR-665-5p | -73.50 | 165 | -32.36 | 232, 532, 400 | 26 | 697 |
|  | miR-709 | -60.91 | 176 | -37.61 | 40, 470 | 19 |  |
|  | miR-6538 | -57.43 | 168 | -35.27 | 432, 399 | 19 |  |
|  | miR-3091-3p | -36.05 | 160 | -18.68 | 403, 535 | 22 |  |
|  | miR-686 | -35.04 | 160 | -19.54 | 204, 116 | 22 |  |
| circ0013866<br>(*Abcc1*) | miR-6902-3p | -39.30 | 159 | -20.30 | 52, 1 | 21 | 441 |
|  | miR-5624-3p | -35.48 | 160 | -20.31 | 415, 163 | 22 |  |
|  | miR-6540-5p | -32.39 | 159 | -16.64 | 414, 163 | 22 |  |
|  | miR-7661-5p | -29.77 | 170 | -18.64 | 168, 209 | 25 |  |
|  | miR-12188-5p | -25.10 | 159 | -25.10 | 201 | 24 |  |
| circ0004838<br>(*Acaca*) | miR-7211-5p | -27.14 | 166 | -27.14 | 290 | 22 | 360 |
|  | miR-7061-3p | -21.73 | 166 | -21.73 | 295 | 22 |  |
|  | miR-181a-2-3p | -22.97 | 159 | -22.97 | 39 | 22 |  |
|  | miR-7094-1-5p | -24.67 | 163 | -24.67 | 334 | 23 |  |
|  | miR-211-5p | -24.02 | 158 | -24.02 | 294 | 22 |  |
| circ0011361<br>(*Ppp2r2a*) | miR-30b-3p | -36.31 | 152 | -20.76 | 358, 597 | 22 | 802 |
|  | miR-6920-5p | -35.33 | 157 | -20.10 | 735, 512 | 22 |  |
|  | miR-1911-3p | -25.61 | 160 | -25.61 | 677 | 22 |  |
|  | miR-6908-5p | -25.52 | 170 | -25.52 | 137 | 22 |  |
|  | miR-693-3p | -24.33 | 168 | -24.33 | 578 | 23 |  |

In the cortex, circ0033491 (*Sez6l*) showed the richest interaction profile, with miR-665-5p exhibiting the strongest binding (total energy −73.5 kcal/mol; three binding sites at positions 232, 532, 400), followed by miR-709 (score 176; two sites) and miR-6538 (−57.4 kcal/mol; two sites). The multiple binding sites per miRNA strengthen the sponge hypothesis and suggest high-avidity sequestration. circ0013866 (*Abcc1*), whose downregulation would release bound miRNAs, showed the strongest predicted binding with miR-6902-3p (−39.3 kcal/mol), miR-5624-3p (−35.5 kcal/mol), and miR-7661-5p (highest score 170), each with two binding positions.

In the hippocampus, circ0004838 (*Acaca*) showed a clustered binding domain between positions 290–334, where four of five predicted miRNA interactions localize. Notably, miR-181a-2-3p, a well-characterized regulator of neuroinflammation and synaptic plasticity, is predicted to be sequestered by the upregulated circ0004838, potentially modulating neuroinflammatory signaling during hippocampal aging. circ0011361 *(Ppp2r2a*), progressively downregulated with age, is predicted to release miR-30b-3p (strongest binding: −36.3 kcal/mol; two sites) and miR-6920-5p (−35.3 kcal/mol; two sites). miR-30b is well-established in synaptic vesicle trafficking and neurodegeneration.

To map downstream regulatory effects, putative miRNA–mRNA interactions were predicted using miRDB (**Supplementary information, S4**). Target genes were ranked according to the miRDB target score, and a score threshold ≥80 was applied to identify high-confidence interactions. For each miRNA, up to 10 target genes were selected based on both prediction of strength and biological relevance to neurodegeneration, synaptic dysfunction, neuronal signaling, and inflammatory pathways. For 2 of the 20 miRNAs, the ≥80 threshold yielded an insufficient number of biologically relevant targets. In these cases, a relaxed score range of 55–70 was applied to incorporate additional genes with known functional relevance to neuronal or neuroinflammatory processes. All selected miRNA–mRNA interactions were integrated into downstream enrichment analyses and incorporated into the circRNA–miRNA–mRNA regulatory network.

The curated circRNA–miRNA and miRNA–mRNA interactions were combined to generate a circRNA-centered ceRNA network. Visualization in Cytoscape revealed a hierarchical structure in which each circRNA served as a regulatory hub connected to its top five miRNAs, which in turn converged on multiple downstream mRNA targets (**Figure 4**).

**Figure 4.**
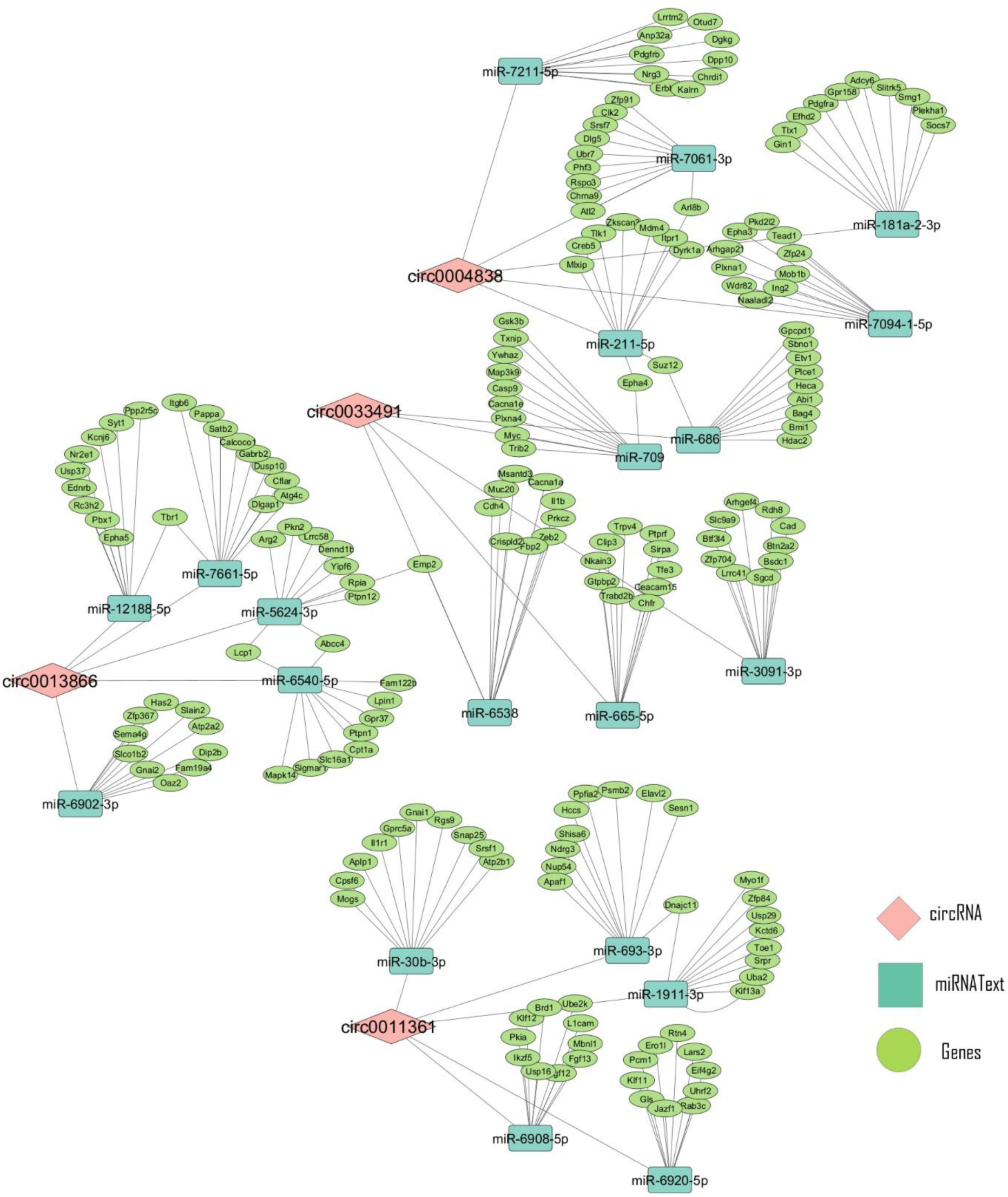
circRNA–miRNA–mRNA regulatory interaction network of key dysregulated circRNAs. Predicted ceRNA (competing endogenous RNA) interaction networks for four consistently dysregulated circRNAs, circ0004838, circ0033491, circ0013866, and circ0011361—in the tauopathy mouse model. Diamond-shaped nodes represent circRNAs, rectangular nodes represent miRNAs, and elliptical nodes represent target mRNAs. Lines connecting nodes indicate predicted sponging interactions. circ0004838 interacts with miR-7211-5p, miR-7061-3p, miR-181a-2-3p, miR-7094-1-5p, miR-211-5p, miR-686, and miR-709, regulating downstream mRNA targets involved in synaptic and transcriptional processes. circ0033491 interacts with miR-7661-5p, miR-12188-5p, miR-5624-3p, miR-6540-5p, and miR-653B. circ0013866 interacts with miR-6902-3p, targeting genes associated with lipid metabolism and signal transduction. circ0011361 interacts with miR-30b-3p, miR-693-3p, miR-1911-3p, miR-6908-5p, and miR-6920-5p, regulating downstream targets implicated in neuronal function and cell cycle regulation. Networks were constructed based on miRNA binding site prediction and expression correlation analysis.

### 3.4. Functional Enrichment Analysis

To elucidate the biological consequences of the identified ceRNA regulatory axes, gene ontology (GO) and Kyoto Encyclopedia of Genes and Genomes (KEGG) pathway enrichment analyses were performed on the mRNA target sets predicted for each circRNA across the biological process (BP), cellular component (CC), and molecular function (MF) domains (**Supplementary information, S5-S8**).

#### Cortical circ0033491 (↑ Upregulation)

Functional enrichment analysis of mRNA targets regulated through the circ0033491/miRNA axis revealed convergent pathways relevant to neuronal apoptosis and synaptic dysfunction. GO CC analysis (**Figure 5a**) highlighted significant enrichment in synaptic and neuronal structural compartments, including the growth cone (n = 4, p = 5.4 × 10⁻³), cell leading edge (n = 3, p = 9.0 × 10⁻³), glutamatergic synapse (n = 6, p = 2.3 × 10⁻²), and neuronal cell body (n = 5, p = 3.1 × 10⁻²), consistent with perturbation of excitatory neurotransmission and axonal guidance machinery. At the MF level (**Figure 5b**), ubiquitin protein ligase binding (n = 6, p = 4.8 × 10⁻⁴) and protein kinase activity (n = 6, p = 4.0 × 10⁻³) were the most enriched terms, indicating that proteasomal substrate recognition and kinase-mediated phosphorylation represent the primary biochemical activities dysregulated by this axis. Calcium channel activity (n = 3, p = 1.9 × 10⁻²) was also significantly enriched, corroborating the cellular component findings. GO BP analysis (**Figure 5c**) identified positive regulation of the apoptotic process as the most significantly enriched term (n = 8, p = 7.0 × 10⁻⁶), alongside negative regulation of insulin receptor signaling (n = 4, p = 1.3 × 10⁻⁴) and negative regulation of neuron projection development (n = 4, p = 7.0 × 10⁻⁴), suggesting that upregulation of circ0033491 drives both neuronal death and impaired structural plasticity. Calcium ions import across the plasma membrane (n = 3, p = 8.2 × 10⁻⁴) and ERK1/ERK2 cascade activation (n = 3, p = 6.1 × 10⁻³) further implicated dysregulated calcium signaling and mitogen-activated kinase pathways in the downstream sequelae. KEGG pathway analysis (**Figure 5d**)identified the insulin signaling pathway (n = 4, p = 7.7 × 10⁻³), MAPK signaling pathway (n = 5, p = 1.1 × 10⁻²), and cell cycle (n = 4, p = 1.1 × 10⁻²) as the most significantly enriched pathways, alongside the Hippo signaling pathway (n = 4, p = 1.1 × 10⁻²) and axon guidance (n = 4, p = 1.5 × 10⁻²). The co-enrichment of insulin resistance (n = 3, p = 4.1 × 10⁻²) and AGE-RAGE signaling (n = 3, p = 3.5 × 10⁻²) further supports a role for metabolic dysregulation and advanced glycation end-product accumulation in circ0033491-mediated tau pathology.

**Figure 5.**
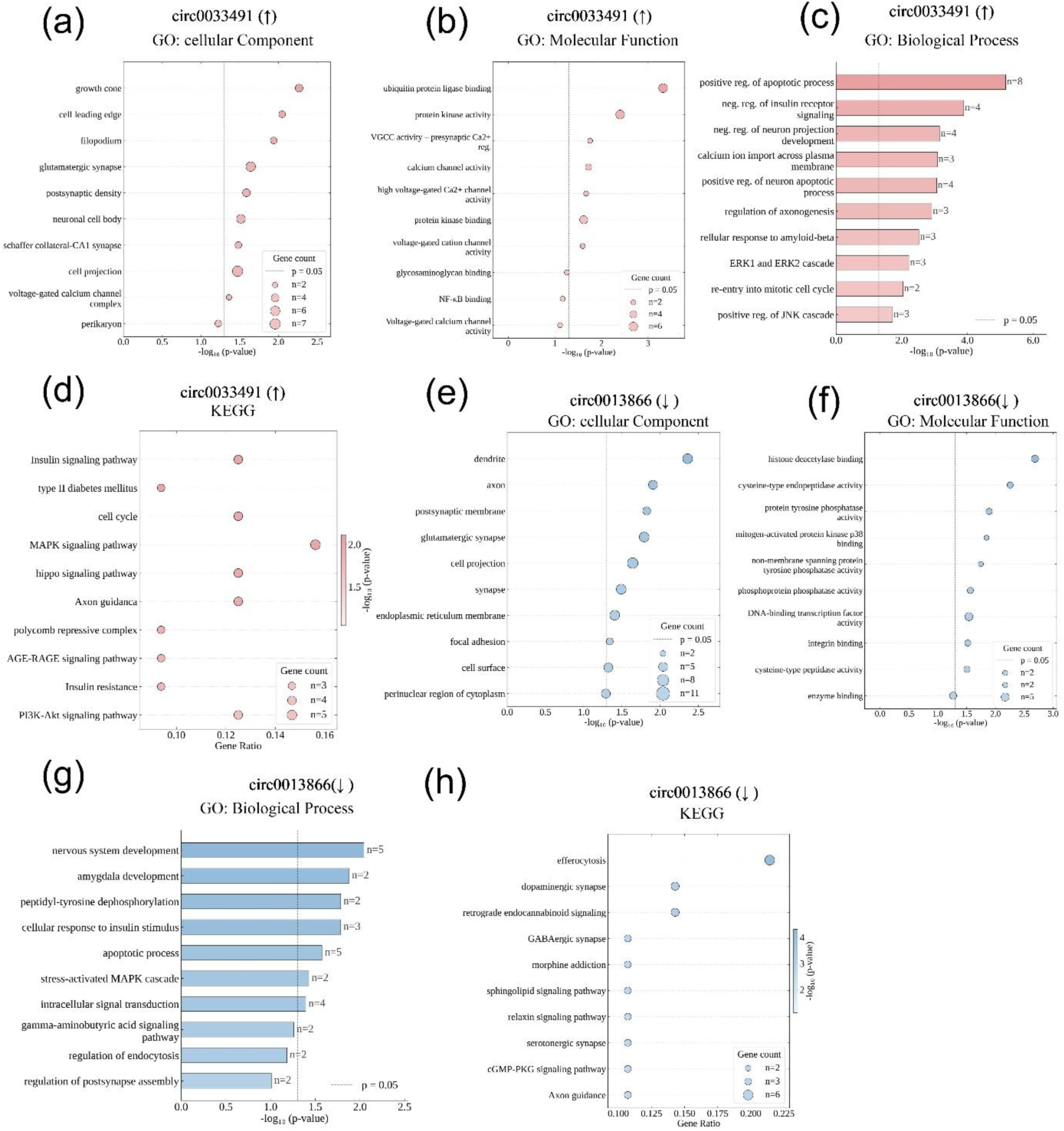
Functional enrichment analysis of circRNA-associated target genes in the cortex of the tauopathy mouse model. (a) GO Cellular Component, (b) GO Molecular Function, (c) GO Biological Process, and (d) KEGG pathway enrichment analysis of upregulated circRNA target genes in the cortex . Dot size reflects gene count and color indicates −log₁₀(p-value). (e) GO Cellular Component and (f) GO Molecular Function (g) GO Biological Process enrichment . (h) KEGG pathway enrichment analysis of downregulated circRNA target genes in the cortex. Dot size reflects gene count; color indicates −log₁₀(p-value). Dashed line indicates p = 0.05 significance threshold.

#### Hippocampal circ0004838 (↑ Upregulation)

Enrichment analysis of circ0004838 target genes revealed a striking predominance of kinase signaling and synaptic organization terms, consistent with a role in disrupting phosphorylation-dependent synaptic homeostasis. GO CC enrichment (**Figure 6a**) was dominated by presynaptic and postsynaptic compartments, with synapse (n = 11, p = 1.3 × 10⁻⁵) representing the most significantly enriched term, followed by glutamatergic synapse (n = 8, p = 8.5 × 10⁻⁴), postsynaptic density membrane (n = 5, p = 1.4 × 10⁻⁴), and postsynaptic density (n = 5, p = 3.2 × 10⁻³). The enrichment of GABA-ergic synapse (n = 4, p = 3.7 × 10⁻³) alongside glutamatergic components suggests that both excitatory and inhibitory synaptic integrity are compromised by this regulatory axis. GO MF analysis(**Figure 6b**) identified kinase activity (n = 12, p = 7.4 × 10⁻⁸) as the most significantly enriched term, with protein tyrosine kinase activity (n = 7, p = 2.3 × 10⁻⁷) and transmembrane receptor protein tyrosine kinase activity (n = 5, p = 2.8 × 10⁻⁶) additionally enriched. The most significantly enriched GO BP terms (**Figure 6c**)were protein autophosphorylation (n = 6, p = 1.8 × 10⁻⁷) and peptidyl-tyrosine phosphorylation (n = 5, p = 2.3 × 10⁻⁷), indicating robust enrichment of receptor tyrosine kinase substrates. Synapse organization (n = 4, p = 5.1 × 10⁻⁴) and synapse assembly (n = 3, p = 7.8 × 10⁻³) were also significantly enriched, implicating structural synaptic remodeling as a downstream consequence of circ0004838 upregulation. KEGG pathway analysis (**Figure 6d**) highlighted gap junction (n = 4, p = 1.0 × 10⁻³) and calcium signaling pathway (n = 4, p = 2.1 × 10⁻²) as the most significantly enriched pathways, with additional enrichment in cholinergic synapse (n = 3, p = 2.9 × 10⁻²) and estrogen signaling pathway (n = 3, p = 3.9 × 10⁻²).

**Figure 6.**
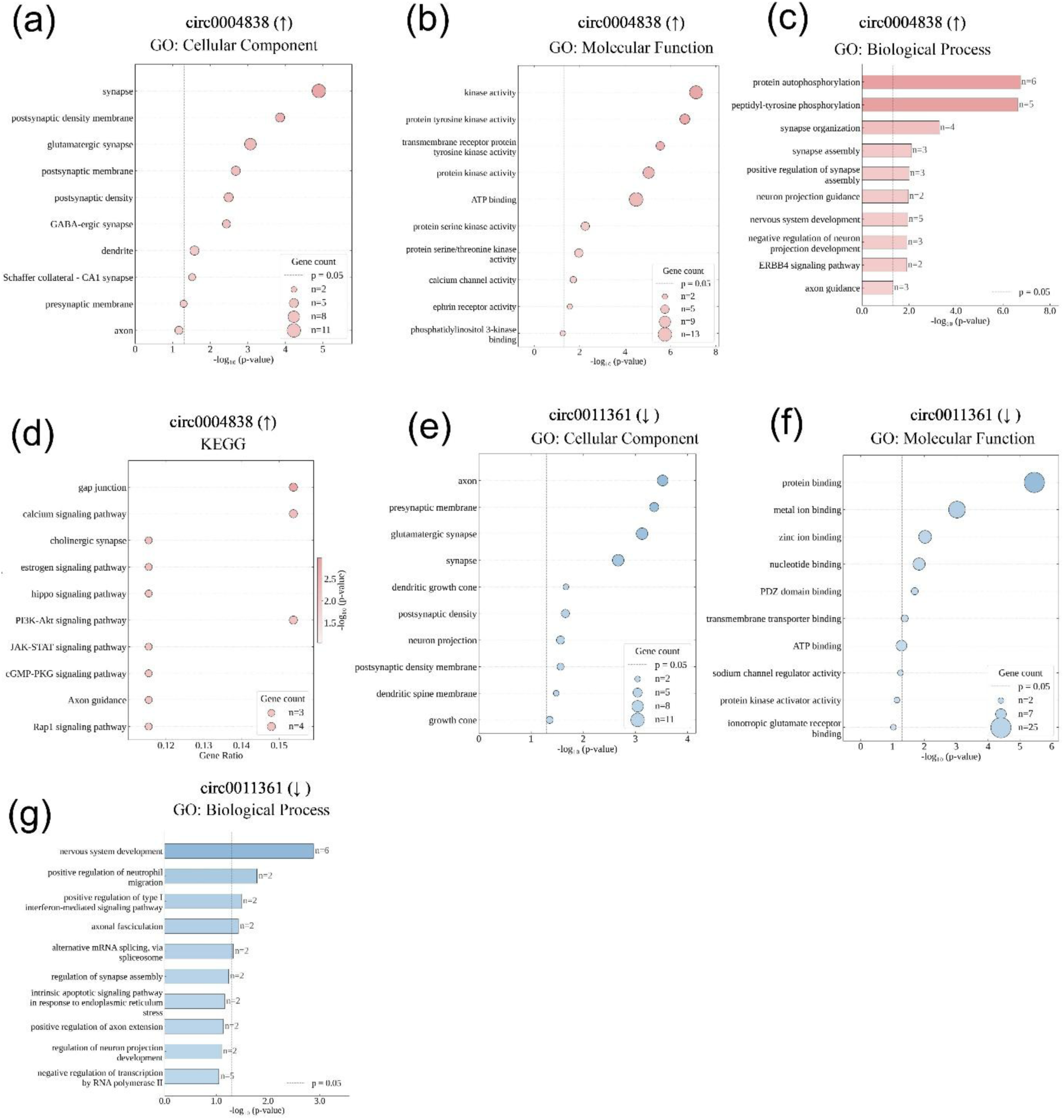
Functional enrichment analysis of circRNA-associated target genes in the hippocampus of the tauopathy mouse model. (a) GO Cellular Component, (b) GO Molecular Function, (c) GO Biological Process, and (d) KEGG pathway enrichment analysis of upregulated circRNA target genes in the hippocampus. (e) GO Cellular Component, (f) GO Molecular Function, and (g) GO Biological Process enrichment analyses of target genes for downregulated hippocampal circRNA . Dot size represents gene count; color indicates −log₁₀(p-value). Dashed line indicates p = 0.05 significance threshold.

#### Cortical circ0013866 (↓ Downregulation)

Loss of circ0013866 expression was associated with enrichment of terms reflecting impaired neuronal survival, synaptic integrity, and phosphatase-mediated dephosphorylation capacity. GO CC analysis (**Figure 5e**) identified the dendrite (n = 6, p = 4.4 × 10⁻³) and axon (n = 5, p = 1.2 × 10⁻²) as the most significantly enriched compartments, with additional enrichment in glutamatergic synapse (n = 6, p = 1.6 × 10⁻²), postsynaptic membrane (n = 4, p = 1.5 × 10⁻²), and endoplasmic reticulum membrane (n = 6, p = 3.9 × 10⁻²). GO MF enrichment (**Figure 5f**) was led by histone deacetylase binding (n = 4, p = 2.1 × 10⁻³), cysteine-type endopeptidase activity (n = 3, p = 5.6 × 10⁻³), and protein tyrosine phosphatase activity (n = 3, p = 1.3 × 10⁻²). GO biological process analysis (**Figure 5g**) identified nervous system development (n = 5, p = 9.0 × 10⁻³) and apoptotic process (n = 5, p = 2.7 × 10⁻²) as the most significantly enriched terms, alongside cellular response to insulin stimulus (n = 3, p = 1.6 × 10⁻²) and intracellular signal transduction (n = 4, p = 4.1 × 10⁻²), indicating that reduced circ0013866 expression compromises neuroprotective insulin signaling and promotes neuronal apoptosis. KEGG pathway analysis identified (**Figure 5h**) efferocytosis (n = 6, p = 4.9 × 10⁻⁵) as the most significantly enriched pathway, suggesting impaired clearance of apoptotic neurons under conditions of reduced circ_0013866 expression. Additional significant pathways included dopaminergic synapse (n = 4, p = 4.9 × 10⁻³), retrograde endocannabinoid signaling (n = 4, p = 6.8 × 10⁻³), GABAergic synapse (n = 3, p = 2.2 × 10⁻²), and morphine addiction (n = 3, p = 2.3 × 10⁻²).

#### Hippocampal circ0011361 (↓ Downregulation)

Functional enrichment of circ0011361 target genes revealed pronounced enrichment in synaptic structural components and protein-binding molecular functions. GO CC enrichment highlighted (**Figure 6e**) the axon (n = 7, p = 3.0 × 10⁻⁴) and glutamatergic synapse (n = 8, p = 7.4 × 10⁻⁴) as the most significantly enriched compartments, followed by synapse (n = 8, p = 2.1 × 10⁻³), presynaptic membrane (n = 5, p = 4.4 × 10⁻⁴), postsynaptic density (n = 4, p = 2.2 × 10⁻²), and dendritic spine membrane (n = 2, p = 3.3 × 10⁻²). GO MF analysis identified (**Figure 6f**) protein binding (n = 25, p = 4.0 × 10⁻⁶) as the most significantly enriched term, with metal ion binding (n = 17, p = 9.4 × 10⁻⁴) and zinc ion binding (n = 10, p = 9.5 × 10⁻³) additionally enriched. GO biological process analysis identified (**Figure 6g**) nervous system development (n = 6, p = 1.3 × 10⁻³) as the most significantly enriched term, alongside axonal fasciculation (n = 2, p = 3.7 × 10⁻²) and positive regulation of neutrophil migration (n = 2, p = 1.6 × 10⁻²). For the selected mRNA of circ0011361, David bioinformatics exhibit no KEGG pathway.

## 4. Discussion

The present study provides the first comprehensive characterization of the circRNA transcriptome across two brain regions and three disease stages in a murine primary tauopathy model. By integrating hierarchical clustering, differential expression analysis, Venn-based persistence filtering, expression trajectory profiling, functional enrichment, circRNA–miRNA thermodynamic binding prediction, and ceRNA network construction, we identify four circRNA candidates, circ0033491/*Sez6l* and circ0013866/*Abcc1* in the cortex, and circ0004838/*Acaca* and circ0011361/*Ppp2r2a* in the hippocampus, whose dysregulation might mechanistically linked to tau hyperphosphorylation, lysosomal tau clearance failure, synaptic structural deterioration, and metabolic dysregulation. A central and unifying finding is that all four circRNA–miRNA–mRNA regulatory axes converge on worsening tau pathology, irrespective of whether the circRNA is upregulated or downregulated, suggesting that the circRNA transcriptome functions as an amplifier rather than a suppressor of tauopathy progression. These findings have significant implications for understanding the molecular architecture of brain aging and tau-related neurodegeneration and support the hypothesis that these circRNA–miRNA axes as good targets for potential therapeutic interventions.

### 4.1. Region-specific dysregulation of circRNA expression

Our findings demonstrate that circRNA expression in tauopathy is region-specific, with the cortex and hippocampus maintaining distinct expression pattern. This observation is consistent with prior transcriptomic studies showing that circRNAs exhibit pronounced spatial specificity in the brain and reflect underlying differences in neuronal subtype composition, circuit architecture, and activity-dependent transcriptional regulations.(Dong et al., 2023; Sekar and Liang, 2019) The clear segregation of cortical and hippocampal samples across all age points indicates that circRNAs encode stable regional molecular identities that persist even as neurodegenerative processes progress. Region-specific circRNA dysregulation likely mirrors well-established patterns of selective vulnerability in tauopathies, where cortical and hippocampal neurons exhibit divergent molecular responses despite shared pathological hallmarks.(Fu et al., 2018) In the cortex, persistently dysregulated circRNAs were functionally linked to glutamatergic synapses, lipid transport, and memory-related processes, pathways known to be disrupted early in tau-mediated cortical dysfunction.(Broekaart et al., 2025; Chung et al., 2021; You et al., 2015)

In contrast, hippocampal circRNA dysregulation was associated with nuclear organization, cytoskeletal remodeling, and metabolic signaling pathways, including AMPK-mediated energy homeostasis.(Dharshini et al., 2019; Wang et al., 2019) This aligns with extensive evidence that the hippocampus is particularly sensitive to metabolic stress and structural plasticity demands during aging and neurodegeneration.(Cai et al., 2026; Kelly et al., 2025)

### 4.2. circRNA-mediated ceRNA regulation in neurodegeneration

Beyond region-specific circRNA expression changes, our integrative analyses reveal that aging-associated circRNAs are embedded within structured ceRNA networks that converge on key molecular pathways implicated in tau pathology. By coupling circRNA expression profiles with predicted miRNA binding and downstream mRNA targets, we demonstrate that circRNA dysregulation has the potential to propagate regulatory effects across multiple biological layers, linking non-coding RNA dynamics to kinase signaling, synaptic integrity, metabolic stability, epigenetic regulation, and neuroinflammatory responses. The cortical circ0033491-centered network represents a prototypical example of how circRNA upregulation may amplify tau-relevant pathology. Circ0033491 is predicted to sequester multiple miRNAs (e.g., miR-709, miR-686, miR-665-5p), whose downstream targets include *Gsk3b, Map3k9, Casp9, Hdac2, Suz12, and Txnip.* These genes collectively span tau kinase activation, MAPK/JNK stress signaling, apoptotic execution, oxidative stress, and epigenetic repression.(Chakraborty et al., 2024; Drewes et al., 1992; Rohn et al., 2002; Sayas and Avila, 2021; Xu et al., 2011) The convergence of these targets within a single circRNA-miRNA axis suggests that circ0033491 may function as an integrative hub reinforcing multiple pro-degenerative processes rather than acting through isolated pathways. Similarly, hippocampal circ0004838 engages miRNAs whose targets include *Dyrk1a, Epha4, Itpr1, Erbb4, Pdgfra, Adcy6*, and *Socs7*, implicating receptor tyrosine kinase signaling, calcium dysregulation, excitatory–inhibitory imbalance, and insulin/AKT/GSK3β coupling.(Carlyle et al., 2014; Egorova and Bezprozvanny, 2018; Majumder and Dutta, 2024; Melchior et al., 2019; Rosenberger et al., 2014; Walker et al., 2015; Woo et al., 2010) The strong enrichment of kinases and phosphorylation-dependent signaling nodes within this network aligns with the central role of aberrant phosphorylation in tau aggregation and supports the notion that circ0004838 upregulation may sensitize hippocampal circuits to tau-driven dysfunction.

In contrast, downregulated circRNAs preferentially map to networks that normally buffer neuronal homeostasis. Cortical circ0013866 targets miRNAs whose downstream genes include *Ppp2r5c, Mapk14, Sigmar1, Slc16a1, Cpt1a, Atg4c, Dlgap1, Gabrb2*, and *Tbr1.* These genes regulate tau dephosphorylation via PP2A, calcium and stress signaling, metabolic substrate delivery, autophagy initiation, and synaptic scaffolding.(Fiock et al., 2020; Lee and Kim, 2017; Li et al., 2025b; Luo et al., 2026; Rasmussen et al., 2017; Sontag and Sontag, 2014; Sun et al., 2020; Trabjerg et al., 2020; Tsai et al., 2015; Yoon et al., 2025) Reduced circ0013866 expression is therefore predicted to derepress miRNAs that collectively weaken phosphatase activity, synaptic stability, and metabolic resilience conditions known to accelerate tau toxicity. A comparable pattern is observed for hippocampal circ0011361, whose downregulation is linked to miRNAs targeting *Snap25, Atp2b1, Ube2k, Psmb2, Mbnl1, Rtn4, Lars2*, and *Shisa6*. These genes are essential for synaptic vesicle release, calcium extrusion, proteostasis, mitochondrial translation, and axonal integrity.(Berrocal and Mata, 2023; Brinkmalm et al., 2014; Carpentier et al., 2014; Jiang et al., 2025; Klaassen et al., 2016; Kulczynska-Przybik et al., 2021; Qian et al., 2024; Song and Jung, 2004)

### 4.3. Disruption of tau phosphorylation homeostasis

Our functional enrichment analysis revealed that the upregulated circRNAs, circ0033491 and circ0004838, drive enrichment of kinase-related GO MF terms including protein kinase activity, protein tyrosine kinase activity, and protein serine/threonine kinase activity, alongside KEGG pathway enrichment in MAPK, insulin and PI3K-Akt signaling. These are pathways that converge on major tau kinases, known to regulate tau phosphorylation.(Deng et al., 2009; Rosenberger et al., 2016; Shomali and Trempe, 2026)

Conversely, the downregulated circRNAs, circ0013866 and circ0011361, are associated with loss of phosphatase-encoding and phosphatase-regulatory gene expression. Protein phosphatase 2A (*Pp2a*) is the dominant tau phosphatase in the human brain, responsible for dephosphorylating tau at the majority of AD-associated epitopes, and its activity is substantially reduced in AD brain tissue relative to age-matched controls.(Gong et al., 1995) The downregulation of circ0013866 thus compromises the *Pp2a* regulatory network precisely when kinase activity is elevated by the upregulated circRNAs, creating a dual insult that simultaneously removes the brake and activates the accelerator of tau phosphorylation.

### 4.4. Synaptic dysfunction as a primary pathological output

A defining feature of the enrichment landscape across all four circRNA target networks is the persistent involvement of synaptic structural and functional components. GO CC enrichment consistently identified glutamatergic synapse, postsynaptic density, presynaptic membrane, dendritic spine membrane, and axon as significantly enriched compartments, while GO BP terms including synapse assembly, synapse organization, and synaptic transmission were enriched across multiple circRNA axes. These findings are directly relevant to the clinical observation that synaptic density rather than amyloid plaque load or neurofibrillary tangle number is the strongest structural correlate of cognitive decline in AD.(Colom-Cadena et al., 2020; DeKosky and Scheff, 1990; Ittner et al., 2010) The identification of GABAergic synapse and GABA-A receptor activity enrichment in the circ0013866 target network is particularly significant, as progressive loss of GABAergic interneurons constitutes a well-documented feature of AD and related tauopathies.(Levenga et al., 2013; Mattson, 2020; Smeralda et al., 2024)

### 4.5. Proteostasis disruption and impaired tau clearance

The enrichment of ubiquitin protein ligase binding and ubiquitin-mediated proteolysis in the circ 0033491 and circ0011361 target networks, together with KEGG enrichment of the proteasome pathway, situates these circRNA axes within the proteostatic collapse increasingly recognized as a defining feature of tauopathy. Tau is primarily degraded by the ubiquitin-proteasome system (UPS), and proteasomal impairment results in the accumulation of soluble and oligomeric tau species that seed aggregation into paired helical filaments and fibrillary tangles.(Chesser et al., 2013; Wang et al., 2025) The heat shock protein 70 co-chaperone Bag4, identified as a target of the circ0033491/miR-709 axis, participates in the decision between proteasomal degradation and autophagy of misfolded tau. BAG4 binds Hsc70 via its BAG domain and modulates chaperone activity; BAG family members serve as key determinants of whether misfolded clients are routed to proteasomal degradation or autophagy, constituting a critical proteostatic triage decision.(Karunanayake and Page, 2021; Minoia et al., 2014)

Similarly, enrichment of autophagy and lysosomal organization terms in the circ0004838 target network reflects disruption of the autophagy-lysosome pathway. Post-mortem analysis of brains from patients with familial AD, corticobasal degeneration, and PSP reveals accumulation of autophagic vesicles, colocalization of hyperphosphorylated tau with LC3, and impaired lysosomal integrity across tauopathies, indicating that autophagosome-lysosome pathway defects are a general feature of tauopathy.(Goniotaki et al., 2025; Piras et al., 2016)

### 4.6. Epigenetic dysregulation as an amplifying mechanism

The enrichment of epigenetic regulatory terms, including histone deacetylation, chromatin remodeling, PRC2 H3K27me3, and histone deacetylase binding across both upregulated and downregulated circRNA axes reveals an epigenetic dimension to ceRNA-mediated tau pathology. *Hdac2*, identified as a target of the circ0033491/miR-686 axis, accumulates in AD neurons and represses expression of synaptic plasticity genes including BDNF, *Glur1*, and *Nr2b*, and its pharmacological inhibition restores cognitive function in mouse models of neurodegeneration.(Guan et al., 2009) *Suz12*, a core subunit of the polycomb repressive complex 2 (PRC2), catalyzes H3K27me3, the repressive histone mark most broadly associated with silencing of neuroprotective gene programmed in ageing and AD.(von Schimmelmann et al., 2016)

### 4.7. Neuroinflammation and the neuroimmune interface

The enrichment of neuroinflammatory pathway components, including TNFα signaling, NF-κB activation through *Zfp91*, interleukin-1β signaling through *Il1β*, and efferocytosis across multiple circRNA target networks connects these ceRNA axes to the neuroinflammation, an established feature of tauopathy.(Vogels et al., 2019; Wang et al., 2022) (Ising et al., 2019; Pampuscenko et al., 2023) The striking enrichment of efferocytosis as the most significantly enriched KEGG pathway in the circ0013866 network reflects impaired phagocytic clearance of apoptotic cells, resulting in an amplification of neuroinflammation.(Chen et al., 2025; Li et al., 2025a; Zhao et al., 2021)

### 4.8. RNA splicing dysregulation and tau isoform imbalance

A particularly disease-relevant finding is the enrichment of RNA splicing and spliceosome components in the circ0004838 and circ 0011361 target networks. The splicing regulators *Clk2, Srsf7, Mbnl1, Srsf1* and *Elavl2* collectively regulate the alternative splicing of tau exon 10, which determines the ratio of four-repeat (4R) to three-repeat (3R) tau isoforms.(Carpentier et al., 2014; Liu and Gong, 2008; Shi et al., 2011) An imbalanced 4R/3R ratio is the defining molecular feature of 4R tauopathies, including PSP and CBD.(Rosler et al., 2019) *Clk2*-mediated phosphorylation of SR proteins including *Srsf7* directly controls exon 10 splicing, and *Clk2* activity changes in tauopathy. Investigations of human postmortem brain tissue from sporadic AD patients reveal that the splicing patterns of tau, *Clk2*, and their regulatory partners are altered in affected brain areas, with amounts of mRNAs of tau isoforms including exon 10 and an inactive form of *Clk2* significantly increased, suggesting that dysregulation of alternative splicing contributes to sporadic AD.(Glatz et al., 2006) (Shi et al., 2011) Furthermore, *Mbnl1* and *Mbnl2* together regulate the alternative splicing of tau pre-mRNA, with both acting as enhancers of tau exon inclusion, and their combined interaction is required to reverse the mis-splicing of tau induced under pathological conditions, demonstrating that loss of *Mbnl* function directly contributes to tau isoform imbalance.(Buchholz and Zempel, 2024; Carpentier et al., 2014)

### 4.9. Dysregulation of the Cortico-hippocampal Circuit: a working model

Based on our findings, we hypothesize that the regional dysregulation of circ 0033491 and circ 0013866 in the cortex, alongside circ 0004838 and circ 0011361 in the hippocampus, drives a coordinated, circuit-level failure where cortical initiation and hippocampal vulnerability mutually reinforce tau pathology (**Figure 7**). Under this proposed model, the upregulation of circ 0033491 and downregulation of circ 0013866 in the cortex could create a state of unopposed tau hyperphosphorylation by simultaneously activating kinase cascades and disabling dephosphorylation, a deficit that correlates with the earliest stages of tangle formation.(Gong et al., 1995) Hyperphosphorylated tau species then propagate anterogradely to the hippocampal CA1 via established trans-synaptic pathways,(Braak and Braak, 1991) where neurons are rendered uniquely susceptible by their own corresponding circRNA dysregulation which favors tau splicing toward the aggregation-prone 4R isoform (Ding et al., 2012; Hartmann et al., 2001; Qian et al., 2011; Qian and Liu, 2014), and altered proteostasis.(Chesser et al., 2013)

**Figure 7.**
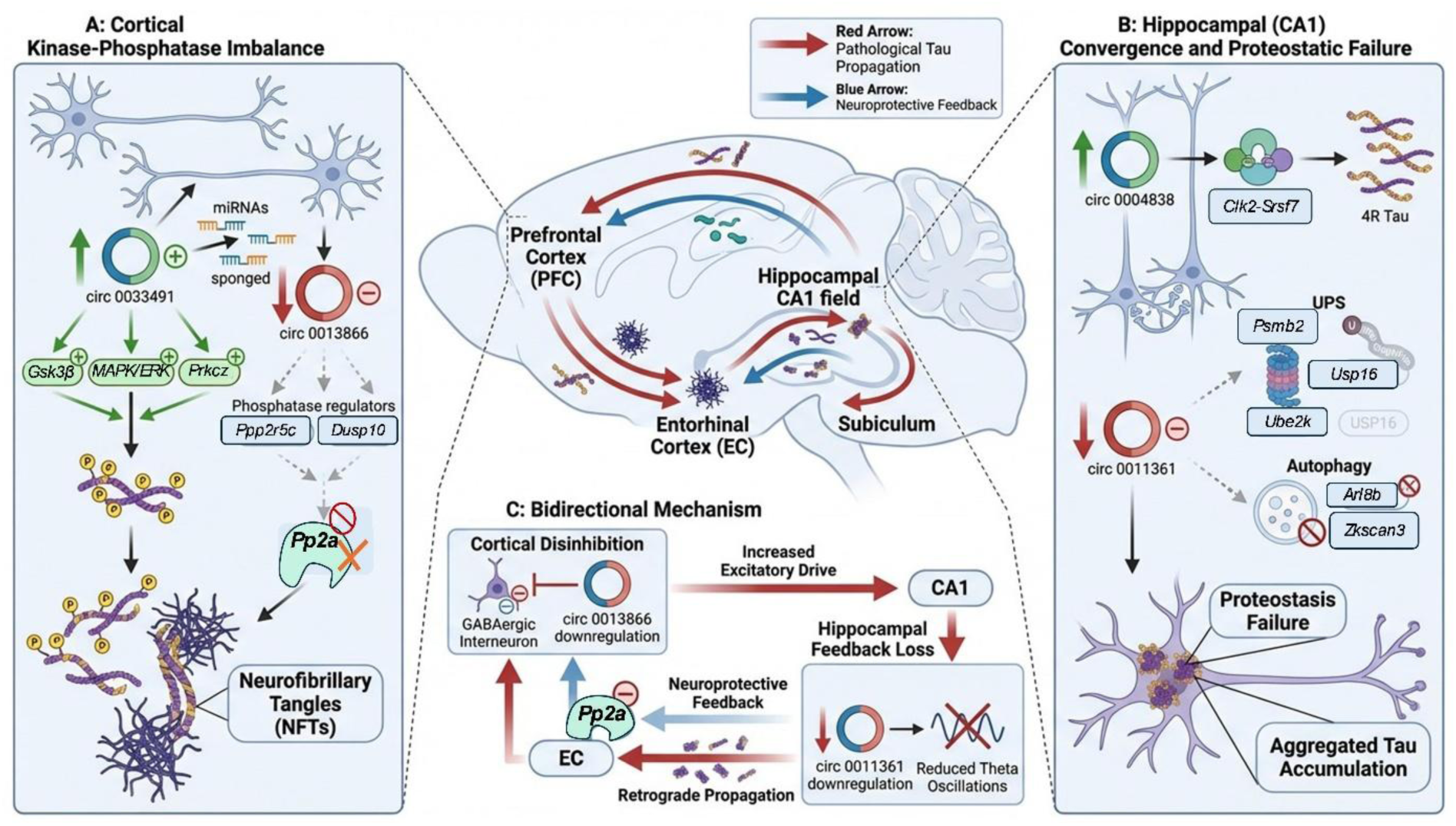
Mechanistic hypothetical model of circRNA-mediated regional vulnerability and bidirectional cortico-hippocampal circuit failure. Central Panel: Schematic of the sagittal mouse brain illustrating the primary anatomical nodes of the cortico-hippocampal circuit (PFC: prefrontal cortex; EC: entorhinal cortex; CA1: hippocampal field; Subiculum). Red arrows indicate anterograde pathological tau propagation, while blue arrows represent neuroprotective feedback and retrograde signals. (A) Cortical Node (Kinase-Phosphatase Imbalance): In the cortex, the upregulation of circ 0033491 "sponges" miRNAs, leading to the release of kinase cascades (*Gsk3β, MAPK/ERK, Prkccz*) from repression. Simultaneously, the downregulation of circ 0013866 leads to the loss of phosphatase regulators (*Ppp2r5c, Dusp10*), resulting in reduced Pp2a activity. This dual dysregulation creates a state of unopposed tau hyperphosphorylation and neurofibrillary tangle (NFT) formation. (B) Hippocampal Node (CA1) (Convergence and Proteostatic Failure): The hippocampal CA1 acts as a convergence point for pathology. Upregulation of circ 0004838 disrupts *Clk2-Srsf7*-mediated splicing, shifting the tau ratio toward the aggregation-prone 4R isoform. Concurrent downregulation of circ 0011361 depletes Ubiquitin-Proteasome System (UPS) components (*Psmb2, Ube2k, Usp16*) and impairs autophagy (*Arl8b, Zkscan3*). This localized proteostatic failure renders CA1 neurons unable to clear arriving cortically derived phospho-tau seeds. (C) Bidirectional Mechanism (Self-Amplifying Circuit Collapse): The circuit failure is non-linear. Cortical Disinhibition: Loss of circ 0013866 in GABAergic interneurons amplifies excitatory drive to the hippocampus. Hippocampal Feedback Loss: Downregulation of circ 0011361 impairs theta oscillations, silencing neuroprotective feedback to the PFC. Retrograde Propagation: Tau oligomers from CA1 propagate to the EC, where they directly suppress Pp2a activity, compounding the cortical phosphatase deficiency. (This figure was generated with the assistance of FigureLab AI)

This pathological circuit is a bidirectional system where cortical disinhibition triggered by the loss of circ 0013866 would increase the excitatory drive to hippocampal synapses,(Hijazi et al., 2020) while the corresponding downregulation of hippocampal circ 0011361 might impair theta oscillations, silencing the neuroprotective feedback required to maintain cortical circRNA stability.(Wang et al., 2021) Our model could explain the clinical dissociation where hippocampal synaptic loss is a more potent predictor of cognitive decline than cortical tangle counts.(Scheff et al., 2007) Therefore, this hypothesis implies that effective therapeutic intervention might not be achievable by targeting a single regional node; instead, it would likely require combinatorial strategies that simultaneously restore cortical kinase-phosphatase balance and hippocampal proteostatic integrity to halt the proposed collapse of the cortico-hippocampal circuit.

### 4.10. Limitations

Several limitations of the present study warrant consideration. The GO and KEGG enrichment analyses are based on predicted miRNA-mRNA target interactions and curated pathway annotations; experimental validation of specific interaction nodes will be required to confirm their functional relevance in neuronal models. The present analysis does not address the temporal sequence of circRNA dysregulation relative to tau pathology progression, a question that longitudinal transcriptomic studies in tauopathy model systems and patient cohorts will be required to resolve. Furthermore, while the ceRNA hypothesis provides a parsimonious framework, circRNAs also function through translation of embedded peptides, interaction with RNA-binding proteins, and modulation of transcription factor activity (Kristensen et al., 2019), mechanisms that may contribute to pathological phenotypes independently of miRNA sponging. Finally, whether the identified circRNA expression changes represent causal drivers of tau pathology or secondary consequences of neurodegeneration remains to be determined through gain- and loss-of-function experiments in vivo.

### 4.11. Conclusions

The present study demonstrates that four circRNAs dysregulated in the context of tau pathology operate as components of functional ceRNA networks that modulate kinase-phosphatase balance, synaptic integrity, proteostasis, epigenetic programming, and neuroinflammation. These findings establish circRNA-mediated ceRNA regulation as a mechanistically tractable and therapeutically actionable layer of post-transcriptional control in tauopathy and identify specific interaction nodes as candidates for future mechanistic and interventional investigations.

## Authors Contributions

Mohammad Shahadat Hossain: conceptualization, experimental design, data analysis, figure preparation and manuscript writing. Domenico Praticò: supervision, conceptualization, experimental design, conceptual guidance, manuscript review and editing.

## Data Availability Statement

The datasets generated and analyzed during this study are available from the corresponding author upon reasonable request.

## Acknowledgements

Dr. Domenico Praticò is the Scott Richards North Star Foundation Chair for Alzheimer research.

## Ethics Statement

All animal procedures were approved by the Institutional Animal Care and Use Committee and conducted in accordance with NIH guidelines for the care and use of laboratory animals.

